# miPIE: NGS-based Prediction of miRNA Using Integrated Evidence

**DOI:** 10.1101/405357

**Authors:** R.J. Peace, M. Sheikh Hassani, J.R. Green

## Abstract

Methods for the de novo identification of microRNA (miRNA) have been developed using a range of sequence-based features. With the increasing availability of next generation sequencing (NGS) transcriptome data, there is a need for miRNA identification that integrates both NGS transcript expression-based patterns as well as advanced genomic sequence-based methods. While miRDeep2 does examine the predicted secondary structure of putative miRNA sequences, it does not leverage many of the sequence-based features used in state-of-the-art de novo methods. Meanwhile, other NGS-based methods, such as miRanalyzer, place an emphasis on sequence-based features without leveraging advanced expression-based features reflecting miRNA biosynthesis. This represents an opportunity to combine the strengths of NGS-based analysis with recent advances in de novo sequence-based miRNA prediction. We here develop a method, microRNA Prediction using Integrated Evidence (miPIE), which integrates both expression-based and sequence-based features to achieve significantly improved miRNA prediction performance. Feature selection identifies the 20 most discriminative features, 3 of which reflect strictly expression-based information. Evaluation using precision-recall curves, for six NGS data sets representing six diverse species, demonstrates substantial improvements in prediction performance compared to miRDeep2 and miRanalyzer. The individual contributions of expression-based and sequence-based features are also examined and we demonstrate that their combination is more effective than either alone.

## 1 Introduction

MicroRNAs (miRNA) are short non-coding RNAs, typically 18-25nts, which modulate post-transcriptional expression of messenger RNA (mRNA) transcripts ^1^. As such, miRNA play a central role in cellular regulation. It has been estimated that 60-90% of all mammalian mRNAs may be targeted by miRNAs ^2^. Through comparative expression analyses and gain- and loss-of-function experiments, it has been shown that miRNA regulate the expression of proteins involved in biological development ^3^, cell differentiation ^4^, apoptosis ^5^, cell cycle control ^6^, stress response ^7^, and disease pathogenesis ^8^. Studies have also shown that miRNA play a role in cellular adaptation to severe environmental stresses such as freezing, dehydration and anoxia ^9–11^. For these reasons, the ability to discover novel miRNA is of great importance.

It is believed that most miRNA share a similar biogenesis mechanism. RNA transcripts known as pri-miRNA contain one or more hairpin structures of approximately 70-120nt in length, known as pre-miRNA. Endonucleases (Drosha and Dicer in animals; DCL1 in plants) process the pri-miRNA in order to form duplexes containing one or more mature miRNA sequences. Ultimately, mature miRNA are incorporated into the RNA-induced silencing complex (miRISC), where a miRNA guides the associated RISC proteins to a targeted mRNA strand. The RISC proteins anneal to the target mRNA and either promote degradation or repress translation of the mRNA ^12^.

Computational miRNA discovery techniques can be broadly categorized as either *de novo* miRNA prediction or ex-pression-based (NGS-based) miRNA prediction ^13^. In *de novo* prediction, putative pre-miRNA sequences which form miRNA-like hairpins are extracted from genomic data sets, and these sequences are classified based on the presence or absence of qualities such as structural stability, sequence motifs typical of miRNA, and structural robustness ^14^. The advantage of *de novo* miRNA prediction is that only genomic sequence is required as input, not transcriptomic data. A disadvantage of such techniques is that they are ignorant of the actual expression of the candidate pre-miRNA region and must therefore consider a far greater number of putative miRNA which may never be expressed. Recent advances in *de novo* sequence-based miRNA prediction have been derived primarily through the application of new pattern classification techniques to the miRNA prediction problem ^15^, and the introduction of new classes of classification features ^16^. However, high class imbalance (1:1000 or higher) within genomic data sets limits the effectiveness of *de novo* classifiers on actual datasets, in spite of high performance often reported on small test data sets with artificially balanced frequencies of positive and negative exemplars ^17^.

In expression-based prediction, data are collected from next-generation sequencing (NGS) experiments. These data represent the sequence and relative abundance of all expressed RNA in a sample, including RNA arising from microRNA (true positives) and other sources including mRNA degradation products and other ncRNA. Predictions of novel miRNA are made from NGS data by seeking patterns of read depth (proxy for transcript abundance in the cell) indicative of processing by Drosha and Dicer endonuclease activity ^18^. These techniques also often examine the strength of the miRNA:miRNA* duplex corresponding to the mature miRNA and miRNA* regions within a putative pre-miRNA region ^19^. Expression-based techniques for miRNA prediction have seen success in recent years ^20–25^, which can be explained in part by the lower class imbalance present in NGS data sets. The number of false positives in a typical NGS experiment is on the order of tens of thousands ^18,26^, whereas one expects tens of millions of miRNA-like structures in a typical genome ^17^Expression-based methods need only evaluate expressed regions, whereas *de novo* methods must evaluate all putative regions capable of forming hairpin structures. Furthermore, methods such as miRDeep2 ^27^ often filter by transcript abundance, considering only the most highly expressed regions as a means to reduce their computational runtime.

Considering both *de novo* and expression-based miRNA prediction techniques, multiple categories of sequence- and expression-based features have been explored, where each may provide independent support for the prediction of miRNA within NGS data sets. State-of-the-art expression-based miRNA prediction techniques, however, only leverage a limited set of these lines of evidence. MiRDeep2^27^ predicts miRNA based on the match between expression levels of NGS read data and expected Dicer processing, the stability of the miRNA:miRNA* duplex, and the significance of the minimum free energy of the pre-miRNA hairpin. MiRanalyzer^28^ predicts miRNA based on secondary structure features, total read depth within the pre-miRNA region, expression of expected Dicer products, and minimum free energy features. These two methods have emerged as standards within the field of expression-based miRNA prediction ^29^. No extant miRNA prediction method employs the full range of features available from NGS data sets, which include both sequence-based and expression-based features. We hypothesize that through the combination of all previously described independent lines of evidence, miRNA prediction performance can be improved.

We here improve on the state-of-the-art performance of expression-based miRNA prediction by integrating the full range of sequence-based and expression-based features to create a novel miRNA predictor. We refer to this new method as miPIE (miRNA Prediction using Integrated Evidence). Our predictor is built using rigorous machine learning techniques, and tested using the metrics of recall and precision, which are directly applicable to real-world miRNA prediction. Additionally, unlike previous methods for expression-based miRNA prediction ^27,28^, all features used in our experiment are invariant to experiment size (total read depth). As NGS technology improves and read depths continue to increase, it is important that all features have this property in order for predictors to be effective on future data sets.

## 2 Methods

### 2.1 Data set selection

Sample data were selected from the NCBI GEO database, using a query consisting of the keywords “ small RNA” and an organism name. Samples were selected for the following criteria: Extracted molecule is “ total RNA”, no infections or knockouts present in the cell, and size fractionation selection is for “ small RNA”. Samples GSE100852 and GSE74879 were collected using the Illumina HiSeq 2500 instrument, sample GSM2095817 was collected using Illumina Genome Analyzer 2, sample GSM1901968 was collected using Illumina HiSeq 1000 and all other samples were collected using the Illumina HiSeq 2000 instrument. Table 1 describes the data sets that were retrieved for this experiment.

**Table 1.**
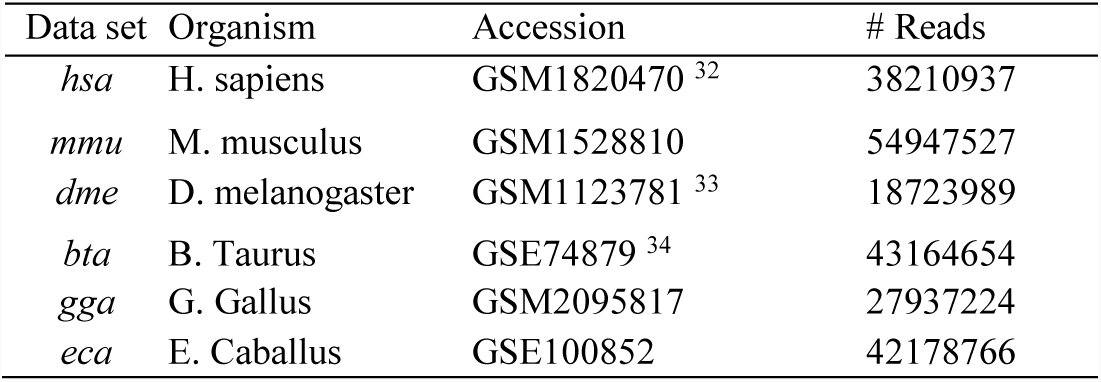
NGS data sets examined in this article

| Data set | Organism | Accession | # Reads |
| --- | --- | --- | --- |
| *hsa* | H. sapiens | GSM1820470 <sup>32</sup> | 38210937 |
| *mmu* | M. musculus | GSM1528810 | 54947527 |
| *dme* | D. melanogaster | GSM1123781 <sup>33</sup> | 18723989 |
| *bta* | B. Taurus | GSE74879 <sup>34</sup> | 43164654 |
| *gga* | G. Gallus | GSM2095817 | 27937224 |
| *eca* | E. Caballus | GSE100852 | 42178766 |

For each of the six samples collected, the following procedure was performed in order to develop positive and negative training sets: The miRDeep2 ^27^ pre-processing algorithm (as implemented in “ mapper.pl”) was applied to all data sets. This tool maps each read stack with at least 4 reads to the reference genome. Putative pre-miRNA regions are extracted (-10/+70nt and -70/+10nt windows based on locally maximal read stacks) and the secondary structure was computed to check for hairpin structures. Running the main script (mirDeep2.pl) resulted in a set of candidate pre-miRNA, each represented by a pre-miRNA sequence, pre-miRNA structure, and the set of reads which map to the sequence (mature, miR-NA*, and loop regions). For each sample, all candidate pre-miRNA which were matched to known high-confidence miRNA from miRBase 21.0 ^35^ using the miRDeep2 “ quantifier.pl” algorithm were selected as true positive samples for training and test.

Each candidate mature miRNA not identified as miRNA in the previous step (labelled as predicted miRNA) was then aligned to the respective species’ coding region data. Alignment was performed using bowtie ^36^. All candidate mature miRNA which aligned to a coding region with at most two mismatches (“ -v 2” bowtie parameter) were selected as negative samples for training and test. Coding region data was retrieved from the Ensembl sequence FTP database ^37^. Table 2 lists the sizes of the final data sets used for this experiment.

**Table 2.** Number of samples in positive and negative classification data sets derived from each NGS experiment data set

| Data set | # of positive genomic regions | # of negative genomic regions |
| --- | --- | --- |
| *hsa* | 167 | 609 |
| *mmu* | 384 | 872 |
| *dme* | 110 | 97 |
| *bta* | 341 | 683 |
| *gga* | 193 | 104 |
| *eca* | 364 | 228 |

### 2.2 Feature set selection

In this study, we examine a set of 223 sequence- and expression-based features. These features incorporate several distinct lines of evidence that have been shown to have predictive power for the classification of miRNA. Of these features, 215 are derived from the feature vector of the sequence-based method HeteroMiRPred ^16^, which in turn gathered these features from a number of methods dating back to 2005. These features all pertain to pre-miRNA sequence and structure and include minimum free energy (MFE)-derived features, sequence/structure triplet features, z-features which encapsulate the significance of the RNA structure relative to those of permuted sequences, and structural robustness features which reflect the ability of the precursor structure to maintain its stability through addition or removal of nucleotides.

Eight expression-based features are added to these sequence-based features. These features are:

(1) Percentage of mature miRNA nts which are paired

(2) Number of pairs in lower stem (outside of mature and miRNA* regions)

(3) Percentage of RNA-seq reads in region which are inconsistent with Dicer processing

(4) Percentage of RNA-seq reads from the loop region which match Dicer processing

(5) Percentage of RNA-seq reads from the mature miRNA which match Dicer processing

(6) Percentage of RNA-seq reads from the miRNA* region which match Dicer processing

(7) Percentage of RNA-seq reads which match Dicer processing

(8) Total number of reads in the precursor region, normalized to experiment size

Here, a match between a read and expected Dicer processing is identical to the definition used by the miRDeep2 study ^27^. A match occurs when a read which maps to a miRNA sequence overlaps the mature, miRNA*, or loop portions of a miRNA with at most 2-nt difference between starting positions on the 5’ ends of the sequences, and at most 5-nt difference between terminating positions on the 3’ ends of the sequences.

These expression-based features provide miRNA classification methods with additional independent lines of evidence for miRNA prediction which are not available from strictly sequence-based feature vectors. Features 1 and 2 provide information on the mature and lower stem regions of the miRNA, while features 3 through 8 provide information regarding the expression pattern within the miRNA region. While the number of expression-based features examined in this study is far less than the number of sequence-based features, it is greater than the number of expression-based features used by both the miRDeep2 scoring algorithm (3: read count within mature region, miRNA* or loop regions; presence of miR-NA* reads matching dicer processing; ratio of reads in the pre-miRNA region which are consistent with Dicer processing) ^27^ and the miRanalyzer scoring algorithm (2: total read count; and expression of miRNA*) ^28^. The remaining features used in these two methods actually pertain to the sequence and/or secondary structure of the putative pre-miRNA or homology to known miRNA.

The final feature set was determined using the correlation-based feature subset selection method ^38^ as implemented in the Weka package ^39^. This algorithm determines an optimal feature set based on correlation between each feature and the class assignment (miRNA vs. pseudo-miRNA), and lack of inter-correlation between features within the feature set. The final feature vector is presented in section 3.1 of this article. Feature selection was performed on the mmu data set, and the resulting feature vector was subsequently employed across all data sets. As a result, performance results for the *mmu* data set represent some optimization using the test set data. This potential source of bias is not evident in the performance results across the *hsa*, *dme*, *bta, gga*, and *eca* data sets where sustained performance is observed.

### 2.3 Classification pipeline

A flowchart illustrating the complete classification pipeline is included in Figure 1. A step-by-step tutorial using sample data for the *dme* species is provided on the website. All miRNA classification in this experiment was performed using a random forest classifier of 500 trees. Trees were built according to the default parameters of the SKLearn random forest library ^40^. Previous studies have demonstrated that random forest classification outperforms competing classifier types for the prediction of miRNA ^17,41^. The miRanalyzer method (see comparison in section 3.2 below) also employs a random forest classifier. Within individual data sets, 10-fold cross validation (10CV) was used to estimate classification performance. Within each fold, the SMOTE algorithm ^42^ was used to oversample the minority class of each training data set to parity with the majority class. Oversampling was performed only on training data sets; class imbalance within each test set was unchanged. When determining performance across data sets, a single classifier was trained using the training data set, and this classifier was used to predict all samples from the hold-out data set.

**Figure 1.**
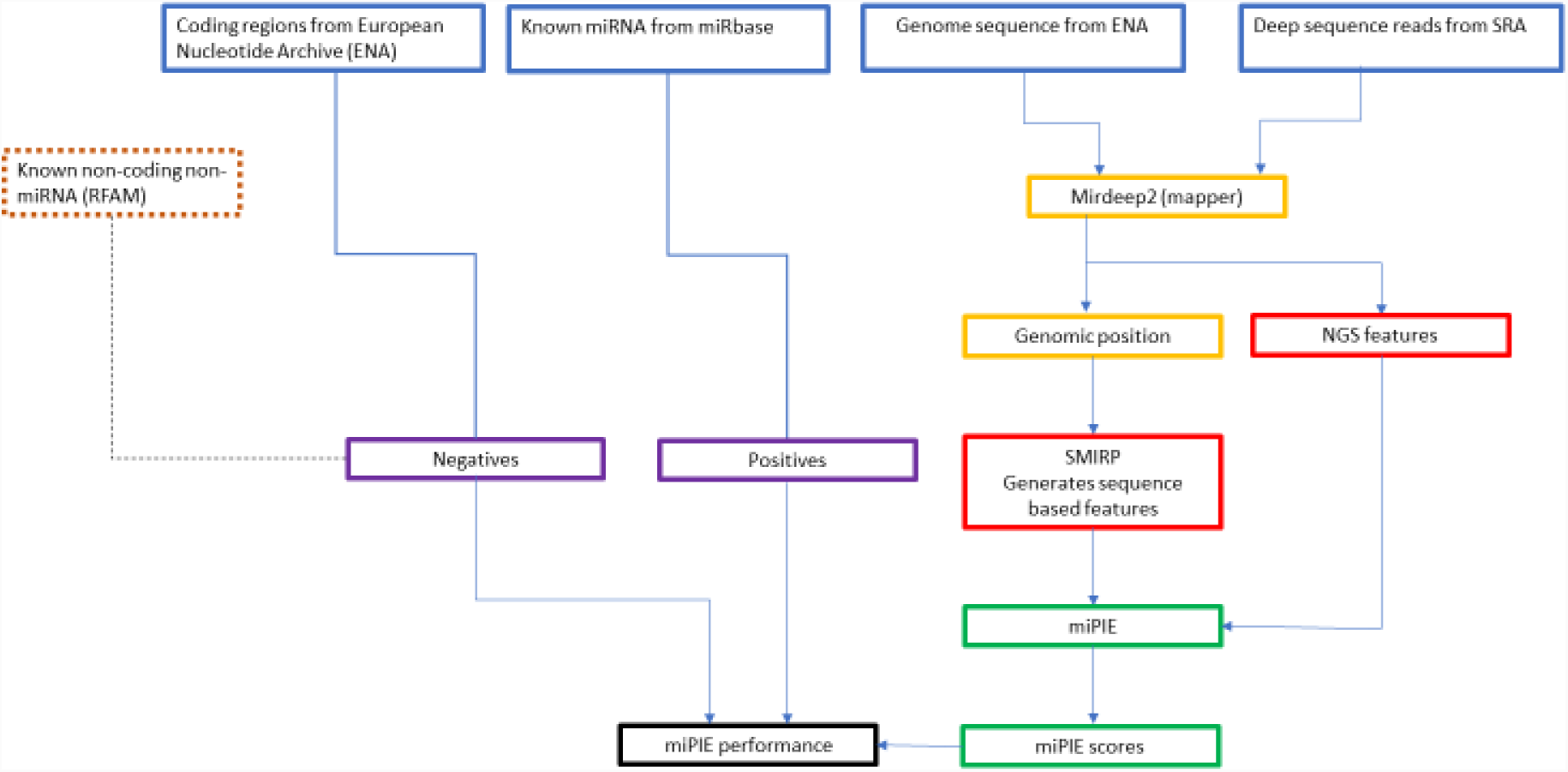
miPIE Pipeline flowchart.

To compute the statistical significance of the observed differences in Re@Pr90 and Re@Pr75 scores between miPIE and miRDeep2 a randomization test was conducted. For the set of ranked results from miPIE and miRDeep2 for a given data set, pseudo samples miPIE* and miRDeep2* were repeatedly developed in the following manner: At each rank, the identities of the patterns at this rank from the miPIE and miRDeep2 result sets were assigned randomly, one to the miPIE* ranked result set and one to the miRDeep2* ranked result set. The differences in Re@Pr75 and Re@Pr90 scores between the two pseudo samples were then computed. In doing so, we enforced the null hypothesis that there is no difference in the way that ranks are assigned by each method. This test was repeated 100,000 times to build a distribution of differences of Re@Pr75 and Re@Pr90 scores expected under the null hypothesis. The percentage of differences which were observed to be equal to or higher than the differences in score between miPIE and miRDeep2 provides the p-value for the observed difference in miPIE and miRDeep2 scores.

### 2.4 Performance metrics

For each data set, the miRDeep2 and miRanalyzer prediction algorithms were run with default parameters over the positive and negative data sets. Performance for all methods is measured using a precision-recall curve (PR-curve). Test set class imbalance is unaltered and represents that of real-world data, as each test set represents the total amount of positive and negative data recovered and processed from an actual NGS experiment. Precision and recall are defined as follows:

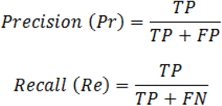

Achievable recall at 75% precision (Re@Pr75) and recall at 90% precision (Re@Pr90) are used as summary statistics. These statistics represent the recall rate achievable at an acceptable degree of precision for experimental validation (75%), and the percentage of miRNA which are correctly classified with very high confidence (90%), respectively.

## 3 Results

### 3.1 Final feature set

The final feature set selected by the correlation feature subset algorithm contains 20 features, which represent seven different classes of evidence for the prediction of miRNA. Sequence-based features relating to secondary structure {MFE3, dH, Tm, Tm/loop}, robustness {SC*absZG, SC/1dp}, base pairing {Probpair 2, 3, 7, 9, 19, and 94}, sequence/structure motifs {“ C((.”, “ T.((“, “ T..(” }, and sequence motifs {“ CG”, “ GA” } were selected. Expression-based features pertaining to miRNA:miRNA* duplex structure (% of paired bases in mature sequence) and Dicer matching of NGS reads (% of reads matching mature, % of reads matching miRNA*) were selected. The fact that automated feature selection arrived at a heterogeneous feature set including both sequence- and expression-based feature supports our hypothesis that an integration of multiple lines of evidence will lead to increased classification performance. By only optimizing the feature set using one of our six datasets, we are implementing a highly conservative validation approach. Optimizing the feature subset over each species will likely lead to further improvements in performance. Descriptions of all features employed by the miPIE classifier are available in Supplemental Table 1.

### 3.2 Performance increase over existing methods

Here we demonstrate the performance increase that our methods achieve over existing state-of-the-art methods for expression-based miRNA prediction. Our method is compared against the miRDeep2 and miRanalyzer methods, over the six data sets described in the methods section. From these six data sets, three of the species (*bta, gga and* eca) have not been previously used by any of the methods.

Figure 2 shows the performance of our method, miRDeep2, and miRanalyzer over the six data sets. While it was possible to create a continuous PR-curve for miRDeep2 by measuring the Pr and Re at various decision thresholds, this was not possible for miRanalyzer since it produces binary predictions without associated probability scores. Therefore, miRanalyzer’ s performance is illustrated as a single point in the P-R space, reflecting its performance at the default decision threshold. As can be seen in these figures, miPIE consistently outperforms both existing methods, particularly for decision thresholds corresponding to high precision (90%).

**Figure 2.**
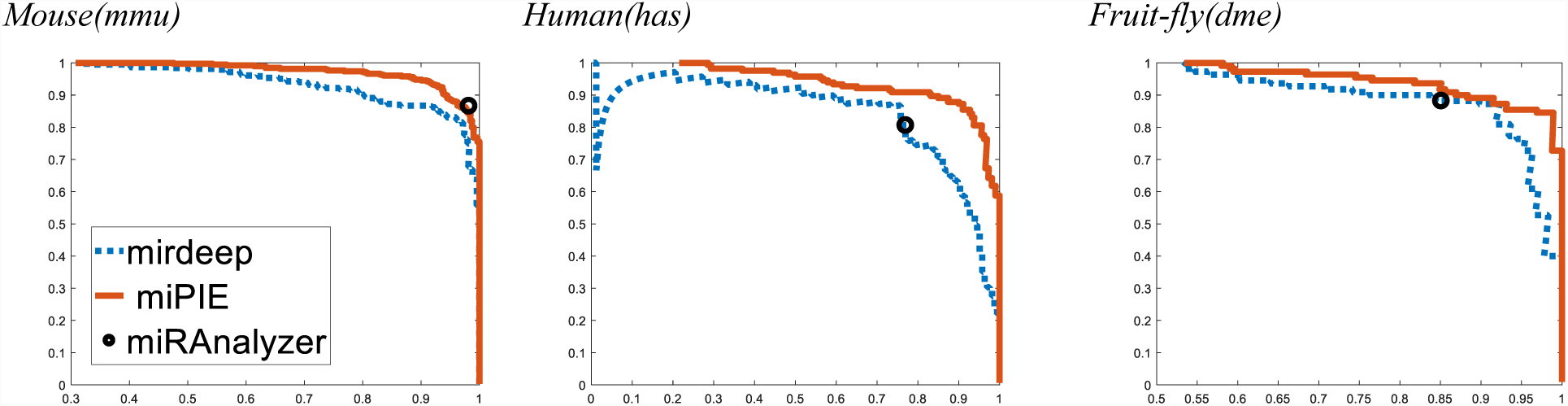
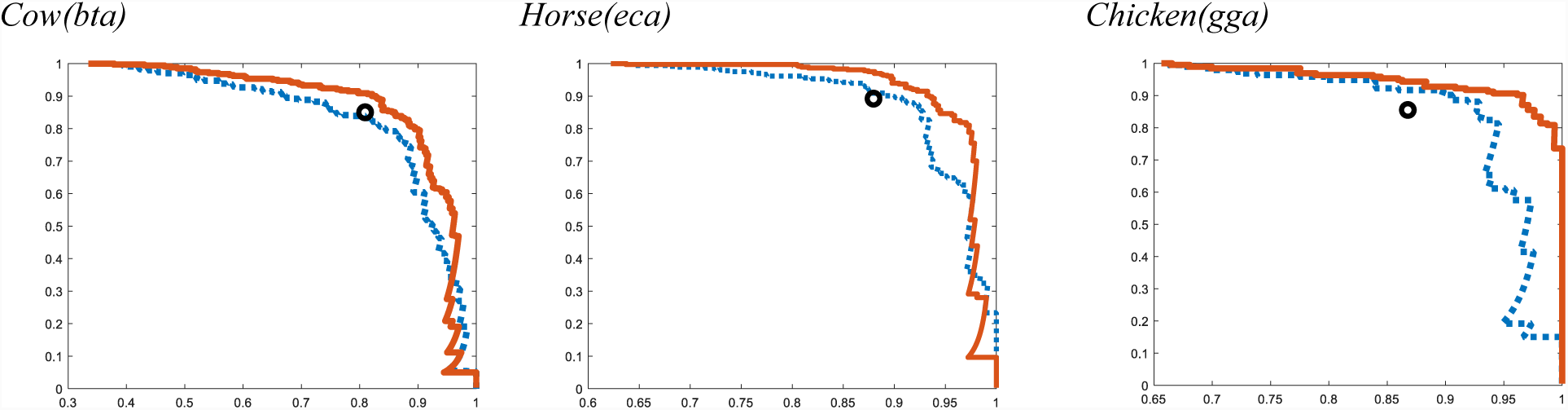
Performance of miPIE, miRDeep2, and miRanalyzer across six data sets. miPIE performance is estimated through 10-fold cross-validation. miRanalyzer produced binary prediction values, so only a single precision level is represented. miPIE outper-forms miRDeep2 and miRanalyzer on all six data sets, with the possible exception of miRanalyzer’ s performance on *mmu*. In all plots, the y-axis represents precision while the x-axis is recall.

Table 3 summarizes the performance increase of miPIE over the miRDeep2 method. On average, our methods increase the number of high-confidence (Pr≥ 90%) miRNA detected by 16%. This improvement is somewhat diminished to 4% at the more permissive decision threshold of Pr≥ 75%. The observed increase in Re@Pr75 is limited by a saturation effect, as both methods are approaching perfect recall at this level of precision. When the randomization test was applied to the six data sets, all of the Re@Pr90 differences were found to be statistically significant (p< 0.01) with the exception of the *hsa* data set. As expected from Figure 2, none of the Re@Pr75 differences was found to be significant at this level, due to the saturation effect discussed above.

**Table 3.** Summary of results comparing miPIE with the state of the art miRDeep2 method, on six NGS data sets. miPIE outperforms miRDeep2 by 16% and 4%, at the 90% and 75% precision thresholds, respectively.

| Data set | Re@Pr75 |  | Re@Pr90 |  |
| --- | --- | --- | --- | --- |
|  | miRDeep2 | miPIE | miRDeep2 | miPIE |
| *mmu* | .930 | .977 (+5.1%) | .867 | .948 (+9.3%) |
| *hsa* | .867 | .909 (+4.8%) | .598 | .873 (+46.0%) |
| *dme* | .909 | .955 (+5.1%) | .873 | .891 (+2.1%) |
| *bta* | .860 | .924 (+7.4%) | .604 | .798 (+32.1%) |
| *gga* | .964 | .985 (+2.2%) | .902 | .927 (+2.8%) |
| *eca* | .975 | .997 (+2.3%) | .897 | .940 (+4.8%) |
| Average | .918 | .958 (+4.0%) | .790 | .896 (+16.0%) |

The original miRDeep2 training data included the three species human (*hsa*), mouse (*mmu*), and fruit-fly (*dme*). Considering this fact, we would have expected a higher performance difference in the three data sets not included in miRDeep2’ s training data. However, no consistent reduction in miRDeep2’ s performance was observed over these new species. The highest performance difference was observed in the human data set. It should be noted that three of these species (*mouse, human, and fruit-fly*) were used to develop both miRDeep2 and miRanalyzer, while the other three (*bta, gga, and eca*) were not used by either. Impressively, performance is largely sustained across the species not used in the development of the two methods (except *bta* for miRDeep2) compared to the three species used.

In order to compare our method with the miRanalyzer method, which provides only binary classification results at a single decision threshold value, Table 4 describes the relative recall rates of miRanalyzer and miPIE at the precision level achieved by the miRanalyzer classifier on each data set. On average, our method increases recall rate by 6.90% relative to miRanalyzer predictions at the stated precision levels. The only dataset on which miRanalyzer performance approaches that of miPIE is *mmu*. This may be explained by apparent similarities between this dataset and that used to train miRanalyzer (NGS series GSE20384), as they both contain mouse testes samples.

**Table 4.** Summary of results comparing miPIE with miRanalyzer using six NGS data sets. When operating at miRanalyzer’ s precision threshold, miPIE outperforms miRanalyzer by 6.9% on average.

| Data set | Precision level | miRanalyzer recall rate | miPIE recall rate |
| --- | --- | --- | --- |
| mmu | .982 | .866 | .841 (-2.9%) |
| hsa | .770 | .806 | .909 (+12.8%) |
| dme | .851 | .882 | .927 (+5.1%) |
| bta | .810 | .849 | .910 (+7.2%) |
| gga | .868 | .854 | .943 (+10.4%) |
| eca | .880 | .891 | .970 (+8.9%) |
| Average | .821 | .840 | .897 (+6.90%) |

### 3.3 Combining sequence- and expression-based features

In order to demonstrate our hypothesis that the predictive power of our method is a result of combining multiple lines of evidence for miRNA prediction, we repeated our 10CV classification pipeline using two subsets of features present in our full original feature set: i) the sequence feature subset containing 20 optimal features selected from the set of all sequence-based features available to our classifier; ii) the expression feature subset, containing the eight expression-based features examined in our study. These feature sets were used to train and test predictors for each of the six data sets described in the methods section. Results of these experiments are shown in Table 5, along with the performance of the miPIE classifier built using the integrated set (i.e. both sequence- and expression-based features). As demonstrated in Table 5, in almost all cases, the performance is improved when sequence and structure features are combined, relative to the use of either class of features alone. In a small number of cases, the combination of features, underperforms one of the individual feature sets alone. The reason for these discrepancies is unclear, however, taken as a whole, these results strongly suggest that the combination of sequence- and expression-based features is preferable to either individual feature set alone.

**Table 5.** Comparison of performance of the integrated miPIE feature set, relative to the performance of similarly trained classifiers trained using only sequence- and only expression-based features.

| Data set | Re@Pr75 |  |  | Re@Pr90 |  |  |
| --- | --- | --- | --- | --- | --- | --- |
|  | all | Seq | Exp | all | Seq | Exp |
| mmu | .977 | .958 | .958 | .948 | .904 | .885 |
| hsa | .915 | .915 | .836 | .879 | .849 | .545 |
| dme | .955 | .982 | .954 | .891 | .918 | .864 |
| bta | .924 | .891 | .891 | .928 | .912 | .969 |
| gga | .993 | .953 | .995 | .798 | .757 | .648 |
| eca | .985 | .953 | .995 | .953 | .929 | .887 |
| Average | .958 | .942 | .938 | .900 | .878 | .800 |

### 3.4 Generalization across experiments

Here we demonstrate the ability of our method to generalize across NGS data sets within and across species. For each of our six data sets, we compare the results of a 10CV experiment (where training and testing data arise from the same NGS experiment) with classification of the same data set using models trained on each of the other five data sets independently, as described in the methods section. Finally, for each hold-out data set, a training set was built using the combination of all other data sets (labeled *all*). Supplemental Figure 1 shows the PR-curves for this experiment. Table 6 summarizes this performance for all six datasets. MiRDeep2’ s performance on the data sets is also shown for comparison. While some decrease in performance is observed when training data arises from a different experiment than the test data, the performance of classifiers trained using all experiments except the test experiment is consistently strong (curve labelled *all* in Supplementary Figure 1).

**Table 6.**
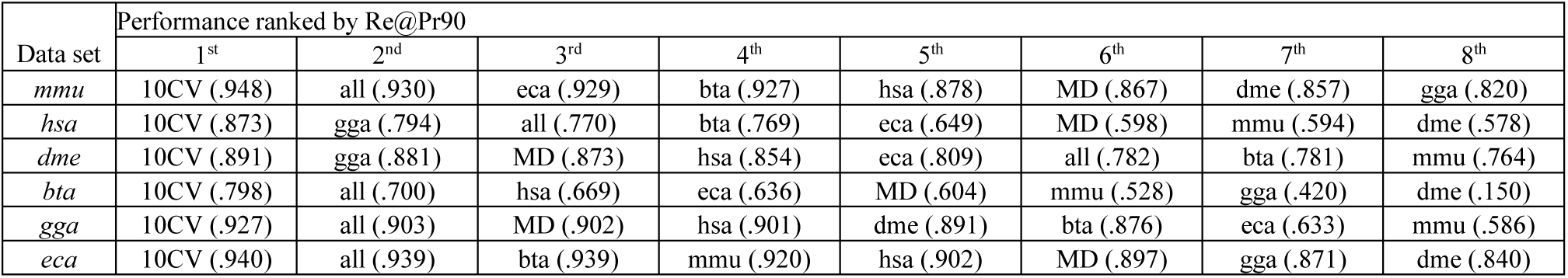
Recall achievable at a precision of at least 90% (Re@Pr90) for 6 test datasets using our method trained over the following datasets: all=combination of 5 datasets, excluding test set; 10CV=10-fold cross-validation over test dataset; hsa = human dataset; bta = cow dataset; mmu = mouse dataset; eca = horse dataset; gga = chicken dataset; dme = fruit-fly dataset. Additionally, miRDeep2’ s performance over each test set is included as MD. Each row is sorted by Re@Pr90 values shown in parentheses.

Table 6 lists the Re@Pr90 for each combined training and test set, in order of decreasing performance for each training set. From these results, we see that miPIE generalizes well to hold-out experimental data sets when data sets from multiple NGS experiments are combined. Our method, when trained using all available data sets except the test species, outperforms miRDeep2 on five of six data sets. Average increase in Re@Pr90 between our method when trained in this manner and the miRDeep2 method is 6.0%. The combined *all* data sets perform as well or better than single experiment data sets in four of six experiments. Fruit-fly is the sole dataset that does not seem to benefit from multi-species pooling of training data. Interestingly, experiments within the same species are not necessarily preferred here (e.g. the top-performing training set for human is the chicken dataset). This result demonstrates that the miPIE method generalizes well to hold-out data sets across experiments and across species when training data is pooled from multiple training experiments.

To demonstrate miPIE’ s ability to predict novel miRNA at high confidence, the final *gga* model was applied to the entire chicken genome. Using a decision threshold of 0.9 resulted in 71 high-confidence novel miRNA predictions (see Supplemental Table 2). Applying a conservative threshold of score levels 10 or 9 (out of a range [-10,10]) to miRDeep2 predictions over the *gga* genome resulted in 46 high-confidence novel miRNA predictions (see Supplemental Table 3). Of the 71 miPIE novel predictions, 27 were uniquely predicted by miPIE (see Supplementary Table 4), while only 2 of the 27 miRDeep2 novel predictions were unique (see Supplemental Table 5). The intersection of miPIE and miRDeep2 predictions results in 44 novel miRNA, each representing novel testable hypotheses (see Supplemental Table 6). A table of all the novel predictions made by miPIE for the 6 test species is presented in Supplemental Table 7.

## 4 Discussion

In this study, we introduce miPIE, a classification method for NGS-based miRNA prediction that integrates both sequence- and expression-based features and employs a rigorous pattern classification approach. All features used by miPIE are independent of the read count of the NGS experiment, such that the performance of this method will remain consistent as NGS technology continues to develop. miPIE is compared with two existing state-of-the-art methods using the metrics of precision and recall, which are directly applicable to the end users of miRNA prediction software in that it answers the two questions: “ Of the actual miRNA in my sample, what percentage will be identified” (recall) and “ Of all the predicted novel miRNA, what percentage will correspond to actual miRNA?” (precision). At high precision levels (90%), miPIE increases recall by 16% relative to the popular miRDeep2 method. Furthermore, miPIE increases recall by 6.9% relative to the miRanalyzer algorithm, at the precision levels reported by miRanalyzer.

One caveat with the 10CV results presented in sections 3.2 and 3.3 is that the 10CV protocol may permit highly similar miRNA (e.g. from same family) to appear in both training and testing subsets. This could lead to overoptimistic evaluation metrics in both the present study and for previously reported methods. However, the cross-species results in section 3.4 appear to allay this concern, as performance is largely sustained when data from different species are used for training and testing of the method.

The negative data used in this study come from protein coding regions, since these regions are believed to exclude miRNA. To confirm that miPIE is effective for arbitrary pseudo-miRNA regions, we added all available non-coding RNA (tRNA, snoRNA, snRNA, rRNA) data from Rfam ^30^ to our negative data sets and re-ran the experiments. Performance was sustained over these broader negative data, confirming that miPIE has not simply learned to recognize protein-coding regions as being negative.

The primary avenue for future improvement of miPIE, and of expression-based miRNA prediction as a whole, is the development of strong training data sets that combine data from multiple NGS experiments. As evidenced in section 3.3 of this study, miPIE’ s generalization performance increases with the incorporation of multiple training data sets, a result which is consistent with that of the miRanalyzer experiment ^28^. Additionally, increasing the quality of training data has proven successful in the field of *de novo* miRNA prediction ^31^. With the ever-increasing availability of NGS data across myriad species, it will be feasible to create larger training data sets incorporating more species. Once training data has been curated from many species, approaches such as used in the species-specific miRNA pipeline ^17^ can be applied to NGS-based miRNA prediction data sets, thereby increasing prediction performance on non-model species. We anticipate that users will gain additional insight by running all three methods (miPIE, miRDeep2, and miRanalyzer). Combination strategies such as the union or intersection of the predicted miRNA will likely increase sensitivity or specificity, respectively. Furthermore, it is suggested that novel features be developed in both the expression and sequence space, and that feature selection be repeated periodically to incorporate the newest and most effective features from both fields. Lastly, miRDeep2’ s decision to examine only the most abundant transcripts will limit the overall recall of the method. With the increase in precision achievable using miPIE, it may be possible to examine a greater number of putative pre-miRNA, while still limiting the expected number of false positive predictions.

## 5 Conclusion

In this paper we introduce a classification method for miRNA prediction by integrating both sequence and expression-based features. Our method uses mirDeep2’ s pre-processing step to identify putative pre-miRNA regions, and performs classification using novel decision logic. By leveraging both expression- and sequence-based features, miPIE increases recall at 90% precision by 16% relative to mirDeep2. In addition, when compared with the miRanalyzer algorithm, an increase of 6.9% in recall is observed at equivalent precision levels. The integrated features used in miPIE are also independent of the NGS experiment read count, ensuring consistency of the method’ s performance with the continuing development of NGS technology. The performance increase observed in miPIE compared to the other well-known methods indicates that integrating sequence and expression-based evidence may enhance our ability in novel miRNA prediction.

## Additional information

### Ethics approval and consent to participate

Not applicable

### Consent for publication

Not applicable

### Availability of data and material

Our method is available as an open source project at http://github.com/jrgreen7/miPIE. All datasets used in this study are available on NCBI-GEO (see Table 1 for accession numbers).

### Authors’ contributions

RP and JG conceived of the study. RP and MSH developed the algorithms and conducted the experiments. RP wrote the initial draft and all authors reviewed and approved of the final manuscript.

#### *Acknowledgements*

The authors would like to thank Kyle K. Biggar for constructive feedback during the development of our method.

